# A New Fluorogenic Substrate for CYP1A2 and Its Application in Studying the Effects of Alcohol Exposure on Liver Drug Metabolism

**DOI:** 10.64898/2026.02.21.703381

**Authors:** Kari Gaither, Nadezhda Davydova, Kannapiran Ponraj, Dilip K. Singh, Bhagwat Prasad, Dmitri R. Davydov

## Abstract

Aiming to develop a high-throughput fluorimetric assay for the activity CYP1A2, we introduced 6-Methoxy-2-naphthoic acid (MONA) as a new fluorogenic substrate for this important metabolizer of antidepressants and psychotropic drugs in human liver. We demonstrated that oxidative demethylation of MONA by liver microsomes results in a red shift and a substantial increase in fluorescence. This effect, which is exceptionally well pronounced at alkaline pH, allowed us to develop a sensitive and robust high-throughput assay of MONA metabolism. Probing the activity of 15 individual recombinant human P450 enzymes, we found that only two P450 species exhibited activity in MONA demethylation: CYP1A2 (*k*_*cat*_=11.9±2.2 min^-1^, *K*_*M*_=578±106 µM) and CYP2A6 (*k*_*cat*_=0.48±0.07 min^-1^, *K*_*M*_=54±15 µM). Since the *K*_*M*_ values of the two enzymes are well resolved and the turnover rate observed with CYP2A6 is much lower than that of CYP1A2, this new fluorogenic substrate is useful as a specific probe for CYP1A2 activity in HLM. Importantly, MONA is not metabolized by CYP1A1 and CYP2C19, which distinguishes it from all known CYP1A2 fluorogenic substrates. We then used MONA to investigate the effects of chronic alcohol exposure on CYP1A2 activity using a series of 23 proteomically characterized individual HLM preparations from donors with various levels of alcohol consumption. The substrate saturation profiles (SSP) acquired with these preparations were subjected to global kinetic analysis by approximating them with combinations of two Michaelis-Menten equations with globally optimized *K*_M_ values of 11 and 553 µM. The amplitudes (*V*_max_ values) of both components showed a pronounced increase with increasing alcohol exposure of the liver donors. The *V*_max_ of the minor high-affinity component was best correlated with the abundance of alcohol-inducible CYP2E1 enzyme. The correlation was further improved by combining it with the abundances of CYP2A6 and CPR. This finding suggests that this minor component reflects the activity of CYP2A6 in the complex with alcohol-inducible CYP2E1 protein. In contrast, the *V*_max_ of the predominant CYP1A2-catalyzed low-affinity component revealed a pronounced correlation with the abundances of CYP1A2 and NADPH cytochrome P450 reductase (CPR). These results suggest a considerable increase in the rate of metabolism of drug substrates of CYP1A2 by chronic alcohol exposure that takes place despite an alcohol-induced decrease in CYP1A2 expression.

## 1. Introduction

Located in the endoplasmic reticulum of liver, lung, intestinal, and other tissues, cytochrome P450 enzymes (P450) are essential to human drug metabolism, as well as the biotransformation of endogenous substrates such as fatty acids and steroids. Understanding P450 activity towards substrates is therefore central to research on drug disposition, drug–drug interactions, and adverse effects. A routine and critical component of understanding drug metabolism and evaluating new drug candidates involves studying the inhibition or activation of P450s by measuring their activity. Commonly used analytical methods such as HPLC, UPLC, and LC-MS/MS are labor-intensive and require extensive sample preparation, costly instrumentation, and specialized personnel, and often involve long run times that limit throughput. Fluorescence-based assays provide an alternative, more efficient, and cost-effective approach to assessing P450 activity. They may be used as a high-throughput tool to study inter-individual variability in P450 metabolism.

CYP1A2 is highly expressed in the liver and is among the five major cytochrome P450 enzymes responsible for most hepatic oxidative drug metabolism, collectively accounting for approximately 80% of clinically relevant drugs [1]. Considerable inter-individual variability in activity and expression has been documented, and additional modulation by environmental and dietary-drug interactions may critically affect CYP1A2-mediated metabolic pathways [2, 3]. A wide variety of substrates are metabolized by CYP1A2, including tricyclic antidepressants and antipsychotics such as amitriptyline, clozapine, haloperidol, and imipramine, many of which have narrow therapeutic windows [2, 3]. Consequently, extensive inter-individual variability could have a significant impact on the pharmacokinetics and clinical outcomes of drugs metabolized by CYP1A2, and as such is an area requiring further study.

Although there are several well-known non-optical probes for CYP1A2, such as caffeine and phenacetin [3, 4], optical probes, such as fluorogenic substrates, are a practical, scalable alternative that is easily adaptable to 96- and 384-well microplate formats. When selective and specific, these probes may be used to explore interindividual variability in P450 activity and to apply them in studies aimed at better understanding drug-drug, herb-drug, and diet-drug interactions.

Several fluorogenic CYP1A2 substrates are currently known [4]. However, all of them are also metabolized by other P450 species, particularly CYP1A1 and CYP2C19. Although 7-methoxyresorufin is considered a specific CYP1A2 probe in liver microsomes, it exhibits similar turnover rates when metabolized by purified CYP1A1 and CYP1A2 [5, 6]. The highly sensitive and high-throughput-friendly probe for CYP1A2 activity, 3-cyano-7-ethoxycoumarin, is also metabolized by CYP1A1, CYP2C19, and CYP2B6 [7, 8]. Therefore, a high-throughput-friendly probe substrate with high selectivity for CYP1A2 would be a notable advancement.

In our recent efforts to develop a specific fluorogenic substrate for CYP4A11 [9], we identified 6-methoxy-2-naphthoic acid, or MONA, as a fluorogenic substrate for CYP1A2. It is worth noting that MONA is a close analog of the non-steroidal anti-inflammatory drug naproxen (6-methoxy-α-methyl-2-naphthaleneacetic acid), which undergoes P450-dependent O-demethylation that also alters its fluorescence. However, O-demethylation of naproxen is catalyzed by several P450 species (CYP2C9, CYP2C8, and CYP1A2), of which CYP2C9 plays a predominant role [10, 11]. In contrast, as we demonstrate in this study, MONA is a highly selective substrate of CYP1A2, which is not metabolized by CYP1A1 or any CYP2C enzyme. We selected it to develop a method to facilitate high-throughput assays for the study of interindividual variability and potential adverse drug interactions within large preparations of HLMs, and to examine further the impact of alcohol exposure on CYP1A2 activity in the human liver.

We describe herein a fluorimetric assay for high-throughput studies of MONA metabolism via O-demethylation, which we employed to investigate the impact of alcohol consumption on the CYP1A2 activity in human liver. Probing 16 recombinant human P450 enzymes with this substrate, we demonstrate that only CYP1A2 and CYP2A6 can metabolize MONA at any appreciable rate. Although recombinant CYP2A6 has a higher affinity to this substrate, the rate of MONA demethylation by this enzyme is negligibly small as compared to that exhibited by CYP1A2. We then assessed MONA metabolism in a set of 23 proteomically characterized microsomal preparations from liver specimens obtained from individuals with varying levels of alcohol exposure. Our results demonstrated a considerable increase in the rate of CYP1A2-dependent MONA demethylation caused by chronic alcohol exposure, despite a significant decrease in CYP1A2 expression, as evidenced in our recent study [12].

## 2. Materials and Methods

### 2.1 Chemicals

6-Methoxy-2-naphthoic acid was purchased from Cayman Chemicals (Ann Arbor, MI, USA). 6-hydroxy-2-napthoic acid was procured from Thermo Fisher Scientific (New Jersey, NJ). Acetoacetanilide was procured from the Tokyo Chemical Industry (Tokyo, Japan). Glucose-6-phosphate dehydrogenase from Leuconostoc mesenteroides was obtained from Worthington Biochemical Corporation (Lakewood, NJ). NADP and glucose-6-phosphate were the products of MilliporeSigma (Burlington, MA, USA). All other reagents were of ACS grade and used without additional purification.

### 2.2 HLM samples from individual donors

In this study, we used 23 microsomal preparations from liver specimens obtained from deidentified individual donors. The set comprises 17 individual liver samples from the biobank established in the Prasad laboratory at WSU Spokane and 6 samples from donors with a history of moderate-to-heavy alcohol consumption procured from BioIVT Corporation (Baltimore, MD). The selection criteria were based on the availability of documented alcohol intake history with varying levels of alcohol consumption, while excluding the use of illicit substances. The group contained 9 HLM samples from non- and light drinkers (<=1 drink per day), 7 samples from moderate drinkers (2-3 drinks per day), and 7 samples from heavy drinkers (>=4 drinks per day). The demographic characteristics of the donors are described in our previous publication [13]. Preparation of the microsomal fraction was performed as described earlier [13].

All HLM preparations were subjected to the determination of relative abundances of the individual cytochrome P450 species, their redox partners (CPR and cytochrome b5), and protein markers of alcohol exposure (HSPA5, PDIA3, P4HB, and CES2 [12]). This analysis was performed using the global proteomics-based total protein approach as described in [13, 14]. Its results are available in the dataset on Mendeley Data [15].

### 2.3. Pooled HLM preparation

In the experiments on the kinetics of MONA demethylation, we used a pooled HLM preparation from 150 donors of mixed gender (INVITROCYP, lot DNJ) purchased from BioIVT Corporation (Westbury, NY). The supplier of the HLM preparations used in our studies, BioIVT Corporation, has declared to adhere to the regulations of the Department of Health and Human Services for the protection of human subjects (45 CFR §46.116 and §46.117) and Good Clinical Practice, (ICH E6) in obtaining the samples of human tissues used for producing the preparations of human subcellular fractions available from these companies.

### 2.4 Microsomes containing recombinant human cytochromes P450

The preparations of insect cell microsomes containing baculovirus-expressed CYP1A1, CYP1A2, CYP2B6, CYP2C8, CYP2C9, CYP2C19, CYP2D6, CYP2E1, CYP3A4, CYP3A5, CYP2J2, CYP4F12, and CYP4A11 enzymes (Supersomes™) were the products of Gentest (Huntsville, AL), now a part of Discovery Life Sciences (Woburn, MA). Bacterial membranes containing human CYP2A6, co-expressed with human CPR (Bactosomes®), were procured from BioIVT Corporation (Baltimore, MD). All those preparations, except for the CYP1A2 and CYP4A11 Supersomes, also contained human cytochrome b5 co-expressed.

### 2.4 Characterization of the content of protein, NADPH-cytochrome P450 reductase, and cytochromes P450 and b5 in HLM

Protein concentrations in microsomal suspensions were determined by the bicinchoninic acid assay. The concentration of CPR in microsomal membranes was determined based on the rate of NADPH-dependent reduction in cytochrome c at 25 °C, and the effective molar concentration of CPR was estimated using the turnover number of 3750 min-1 [8]. The total concentration of cytochromes P450 in HLMs was determined with a variant of the ‘oxidized CO versus reduced CO difference spectrum’ method described earlier [8].

### 2.5. Protein expression and purification

N-terminally truncated (Δ3-20) and C-terminally His-tagged CYP2E1 [16] was expressed in *E. coli* TOPP3 cells and purified as described earlier [8]. The plasmid for expression of N-terminally truncated (Δ2-23) and C-terminally His-tagged CYP2A6 [17] was kindly provided by Emily E. Scott (University of Michigan, Ann Arbor, MI). The protein was expressed in *E. coli* Topp3 cells and purified as described [18].

### 2.6. Incorporation of purified CYP2E1 and CYP2A6 proteins into Supersomes

Incorporation of purified CYP2E1 and CYP2A6 into Supersomes was performed by incubation of suspensions of Supersomes (5 mg/ml protein) in 125 mM K-Phosphate buffer containing 0.25M Sucrose with purified CYP2E1 or CYP2A6 for 12 - 16 hours at 4°C under argon atmosphere at continuous stirring. For the preparation of CYP2A6-containing Supersomes, we used “control” Supersomes containing human CPR and cytochrome *b*_5_ (Discovery Life Sciences, cat. #456244). P450 protein was added in a 2:1 molar ratio to the concentration of CPR determined as described above (section 2.4). In the case of incorporation of CYP2E1 into CYP1A2 or CYP2A6-containing Supersomes, it was added in a 1:1 molar ratio to the P450 content in Supersomes. Following incubation, the suspension was diluted 4 times with 125 mM K-Phosphate buffer, pH 7.4, containing 0.25 M sucrose, and centrifuged at 53,000 rpm (150,000 g) in an Optima TLX ultracentrifuge (Beckman Coulter Inc., Brea, CA, USA) with a TLA100.3 rotor for 90 min at 4 °C. The pellet was resuspended in the same buffer to a protein concentration of ∼5 mg/ml. The amount of incorporated cytochrome P450 was calculated from the difference between the amount of the heme protein added to the incubation media and the amount of the heme protein found in the supernatant. According to this assay, our procedure resulted in ∼90% of the added protein being incorporated into the microsomal membrane.

### 2.5 Fluorescence spectroscopy

Spectra of emission and excitation of fluorescence were acquired with a Cary Eclipse fluorometer (Agilent Technologies, Santa Clara, CA, USA) equipped with either a thermostatted cell holder and motorless magnetic stirrer (Variomag-USA, Port Orange, FL, USA) or a fluorescence plate reader accessory.

### 2.6 Monitoring MONA demethylation by the formation of formaldehyde

Assays of MONA demethylation, monitoring the accumulation of formed formaldehyde, were performed using a high-throughput fluorimetric assay based on the Hantzsch reaction with acetoacetanilide. The assays were executed as described earlier [19].

### 2.7 Fluorimetric assays of MONA metabolism

The process of developing this procedure is outlined in ‘Probing HLM activity in MONA demethylation and developing a fluorimetric method for real-time monitoring’ under Results. The final protocol, which uses an OT-2 liquid-handling robot (Opentrons Inc., Brooklyn, NY) and a Cary Eclipse fluorometer equipped with a plate reader accessory (Agilent Technologies, Santa Clara, CA, USA), is described below.

MONA metabolism experiments were conducted using 12 MONA stock solutions in the Incubation Buffer (0.1 M Hepes buffer, pH 7.4, containing 60 mM KCl), with concentrations ranging from 0 to 4.85 mM, prepared by serial dilution. A 150 mM stock solution of MONA in DMSO was diluted by the Incubation Buffer to prepare the most concentrated (4.85 mM) solution in the series. This series of stock solutions yielded MONA concentrations in the incubation mixture decreasing from 1212 to 0 µM with a dilution factor of 1.73333 (≈√3). Preparation of the multiwell plates and activity assays was carried out according to the procedures described for the MONACRA substrate activity assays. Briefly, each well of the 96-well plate contained 40 µL of the mixture containing 200 µM NADP, 1 µL/mL Glucose-6-phosphate dehydrogenase, and various concentrations of substrate in the Incubation Buffer. After 8 min of preincubation in the heater-shaker module set to 37 °C and 1200 rpm, when the temperature in the wells stabilizes at 29.5–31 °C, the reaction was started by adding 20 µl of microsomal suspension. Incubation of the samples in the heater-shaker continued for 30 min, after which the reaction was quenched by adding 32 µl of 30 mM trichloroacetic acid per well. The assay protocol ends with adding 48 µl of 2.4 M Na-glycine buffer, pH 10.4, to each well. The incubation plate was then centrifuged at 3400 rpm in a Beckman Allegra 6R centrifuge with a GH-3.8 swing-out rotor with multiwell-plate adapters. After 15 min of centrifugation, the plate was scanned with a Cary Eclipse fluorometer with a multiwell-plate accessory. We acquired the spectra of fluorescence emission in the 416–650 nm region with excitation at 375 nm and the slits set at 20-nm bandwidth.

### 2.8 Global kinetic analysis of substrate saturation profiles

The datasets for global kinetic analysis were assembled as the sets of averages of 2–4 SSPs obtained with each HLM preparation. The combined dataset for all twenty-three HLM preparations was subjected to Principal Component Analysis (PCA). The results of PCA were used to find a set of two Michaelis–Menten equations whose linear combinations best approximate each of the individual SSPs as described earlier [13, 20]. To find the parameters of these components, we used a Nelder-Mead optimization procedure combined with the SURFIT algorithm for multidimensional linear regression applied to the first two principal component vectors as described earlier [13]. The optimized set of two kinetic components resolved thereby was used to determine the amplitudes of the two individual phases in each SSP using the Target Transform Factor Analysis (TTFA) technique [21, 22]. The square correlation coefficients for the resulting approximations of the SSPs were always >0.98 and, in most cases, >0.995. The amplitudes of the individual phases of SSPs were then analyzed to probe their correlations with the composition of the cytochrome P450 ensemble in HLM samples. All manipulations with the dataset, PCA, and regression analysis were performed using our SpectraLab software [23, 24] available for download at http://cyp3a4.chem.wsu.edu/spectralab.html.

## 3. Results

### 3.1 Probing MONA demethylation by HLM

To probe the ability of HLM to demethylate 6-Methoxy-2-naphthoic acid (MONA), we used our fluorimetric assay to measure the amount of formaldehyde produced. As shown in Fig 1a, incubation of a pooled HLM preparation with MONA in the presence of a NADPH-generating system results in the accumulation of formaldehyde, with the rate remaining constant for at least 35 min.

**Figure 1.**
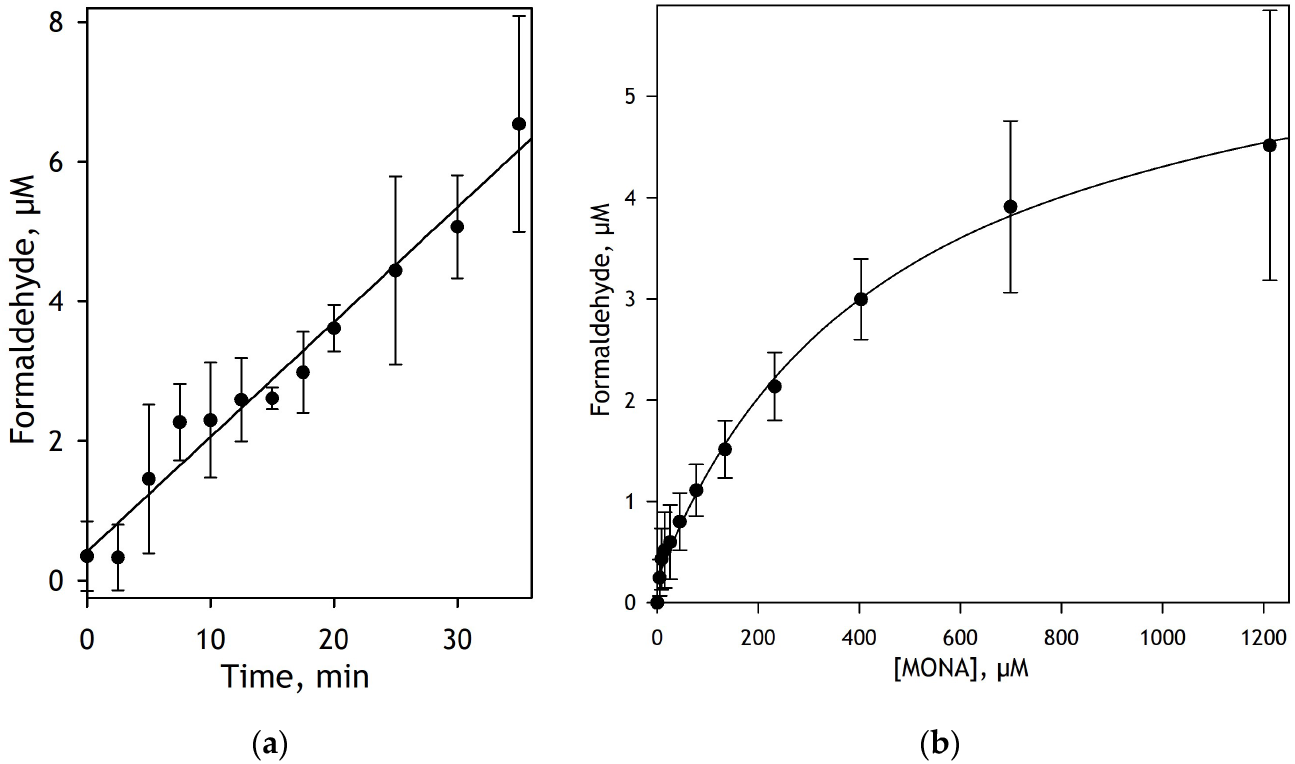
O-demethylation of MONA by pooled HLM monitored by the accumulation of formed formaldehyde. Panel (**a**) shows the kinetics of formaldehyde accumulation in the process of MONA demethylation at 750µM substrate. The dataset represents the average of two individual experiments, and the error bars correspond to the standard deviations. Panel (***b***) shows the substrate dependence of the amount of formaldehyde produced during 30 min of incubation. The dataset represents the average of three individual experiments, and the error bars correspond to the standard deviations. The solid line represents the approximation of the dataset with the Michaelis-Menten equation with *K*_M_=466 µM. Conditions: 750 µM MONA, 1.25 mg/ml HLM, 0.1 M Na-Hepes buffer pH 7.4, 60mM KCl, NADPH-generating system, 30 °C.

The dependence of MONA demethylation rate on substrate concentration is shown in Fig 1b. In this plot, the reaction rate is expressed as moles of formaldehyde formed per minute per mole of microsomal CPR. As seen in this plot, this dependence follows a Michaelis-Menten equation with *K*_M_ = 466 µM. To further study the MONA demethylation process, we developed a fluorimetric assay to monitor it.

### 3.2 Developing a fluorimetric method for monitoring MONA demethylation

To develop a fluorimetric method for monitoring MONA O-demethylation, we studied the fluorescence spectra of MONA and the product of its O-demethylation, 6-hydroxy-2-napthoic acid (HONA). As shown in Fig. 2a, MONA demethylation results in a red shift and a pronounced increase in fluorescence intensity, whereas the excitation band position remains unaffected when measured at neutral pH. These changes can be used in MONA metabolism assays by real-time monitoring of fluorescence increase at 456 nm with excitation at 300 nm. However, the high-throughput acquisition of substrate saturation profiles (SSP) with fluorescence monitoring at neutral pH is complicated by the overlap of the product and substrate emission bands.

**Figure 2.**
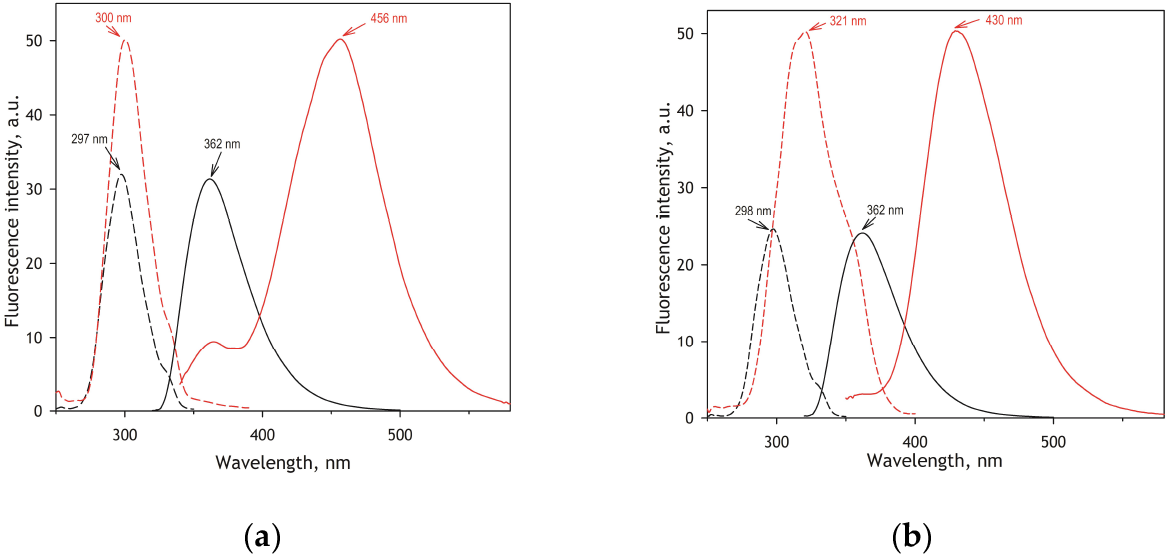
Spectra of fluorescence of MONA (black) and HONA (red) recorded at pH 7.4 (a) and 10.4 (b). Spectra of excitation and emission are shown in dashed and solid lines, respectively.

With this in mind, we explored the effect of alkalizing the assay medium on MONA and HONA fluorescence. The differences in the intensities and positions of maxima of fluorescence between naphthalene methoxy-derivatives with acidic substituent groups and their demethylated products may increase when measured at alkaline pH, as evidenced in the case of naproxen [25] and 3-(6-methoxynaphthalen-2-yl)acrylic acid (MONACRA), a CYP4A11-specific substrate introduced in our recent study [9].

As illustrated in Fig. 2b, alkalization results in a significant red shift of the position of the HONA excitation band, whereas the intensity of MONA fluorescence decreases, with the shape and position of the bands unaffected. This pH effect allows avoiding overlap of HONA fluorescence with that of the substrate when excitation is performed at the red shoulder of the excitation band. We determined that an excitation wavelength of 352 nm provides the best sensitivity for detecting HONA formation, with no interference with the substrate’s fluorescence.

Based on these findings, we developed an automated multiwell-plate-based assay of MONA demethylation. Similar to the setup used with MONACRA, a naphthalene-based CYP4A11 substrate [9], the reaction mixture incubated in the wells is first quenched with trichloroacetic acid and then alkalized by adding concentrated Na-glycine buffer, pH 10.4 (see Materials and Methods).

Application of our method to monitoring MONA metabolism using a pooled HLM preparation is illustrated in Fig. 3. As shown in Fig. 3A, incubating MONA with HLM in the presence of an NADPH-generating system results in a substantial increase in fluorescence, indicating the accumulation of the product of MONA demethylation (HONA). The dependence of the concentration of the formed HONA versus the substrate concentration obeys the Michaelis-Menten equation with *K*_M_=380 µM and *V*_max_=2.4 min^-1^ (Fig. 3B).

**Figure 3.**
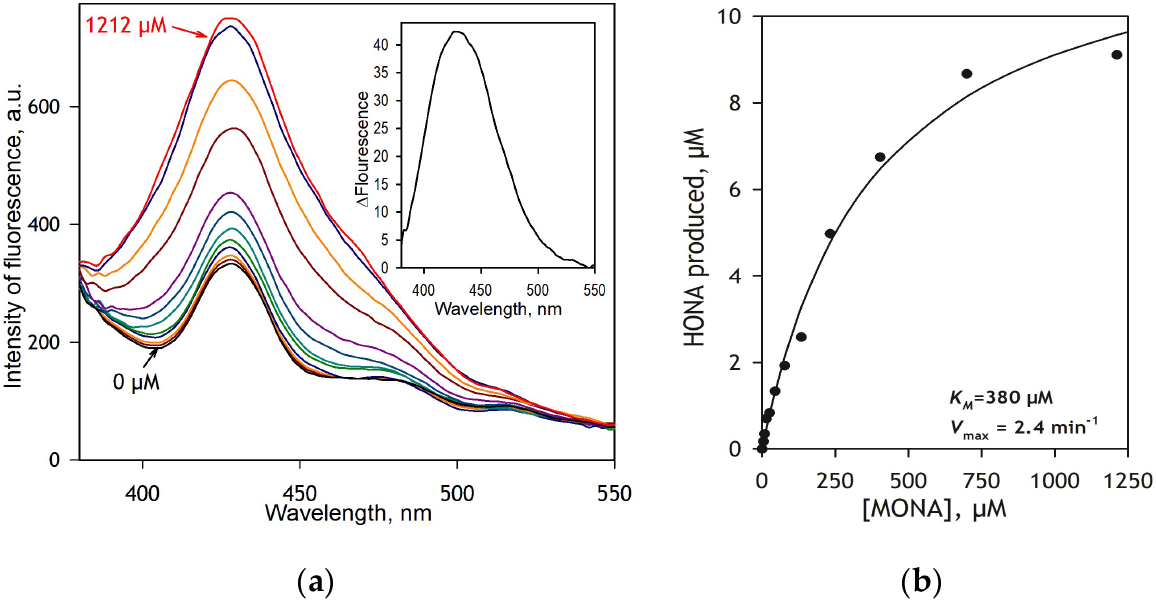
O-demethylation of MONA by pooled HLM preparation monitored by changes in fluorescence. Panel **(a)** shows a series of emission spectra recorded at pH 10.4 with excitation at 356 nm obtained after incubation of HLM (2.5 mg/ml) for 20 min at 30 °C with increasing concentrations of MONA. The inset shows the spectrum of the first principal component obtained by applying PCA to this dataset. A dependence of the concentration of formed HONA on the substrate concentration is shown in panel **(b)**, where the solid line shows the results of fitting the dataset to the Michaelis-Menten equation. The *V*_max_ value given in the plot is calculated per molar concentration of CPR in the incubation mixture.

### 3.2. Activity of recombinant cytochromes P450 in MONA demethylation

We used the high-throughput activity assay described above to probe MONA metabolism by 14 human microsomal P450 enzymes: CYP1A1, CYP1A2, CYP2A6, CYP2B6, CYP2C8, CYP2C9, CYP2C19, CYP2D6, CYP2E1, CYP2J2, CYP3A4, CYP3A5, CYP4A11, and CYP4F12. In these experiments, we used human P450 enzymes co-expressed with NADPH-cytochrome P450 reductase in microsomes of insect cells transfected with recombinant baculovirus (Supersomes®), except for CYP2A6, in which case we used Bactosomes®, where cytochromes P450 were co-expressed with CPR in *E. coli* membranes.

In the case of CYP2A6, we also used a preparation of Supersomes containing CPR and cytochrome *b*_5_, but no P450 enzyme co-expressed (“control” Supersomes), in which CYP2A6 was introduced by incorporating purified protein (see Materials and Methods). All those preparations, except for the CYP1A2 Supersomes and CYP4A11 and CYP2A6 Bactosomes, also contained human cytochrome *b*_5_ co-expressed.

Among all probed P450 enzymes, detectable activity in MONA metabolism was observed with CYP1A2 and CYP2A6 only. The substrate saturation profiles of MONA demethylation by these enzymes are exemplified in Figure 4. Their kinetic parameters are shown in Table 1. As seen from this table, the turnover rate exhibited by CYP1A2 is by far higher than that observed with CYP2A6. However, CYP2A6, while inefficient in MONA demethylation, exhibited higher affinity to this substrate compared to CYP1A2. The difference in the turnover rates between CYP2A6 Bactosomes and CYP2A6-incorporated Supersomes may be, at least in part, explained by the presence of cytochrome *b*_5_ in the control Supersomes used for CYP2A6 incorporation.

**Table 1.**
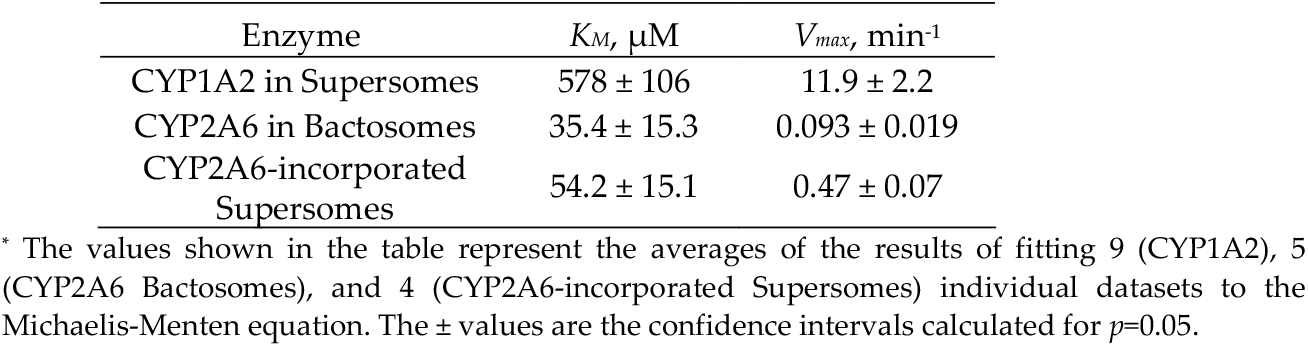
Parameters of MONA demethylation by recombinant CYP1A2 and CYP2A6 enzymes.*

**Figure 4.**
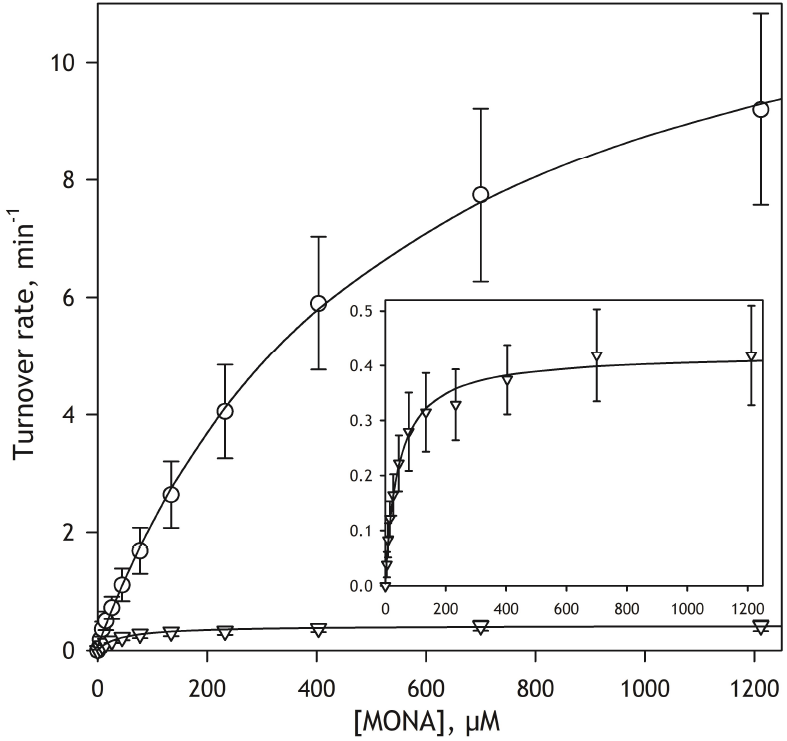
Substrate saturation profiles of MONA metabolism by recombinant CYP1A2 (circles) and CYP2A6 (triangles) enzymes in Supersomes. The inset shows the CYP2A6 dataset zoomed in. Reaction rate is expressed as mol of product per mol of P450 per minute (min^-1^). The datasets represent the averages of 11 (CYP1A2) and 6 (CYP2A6) individual experiments. Error bars show the confidence intervals calculated for *p*=0.05.

### 3.3. Metabolism of MONA by individual HLM preparations and the effect of alcohol consumption

In this study, we capitalized on a collection of HLM samples from donors with varying degrees of alcohol exposure assembled in our laboratory and previously characterized using proteomics [12, 13]. To explore the effects of alcohol-induced changes in the composition of the P450 on the function of CYP1A2, we constructed a set of SSPs in a series of 23 individual HLM preparations, in which the composition of the microsomal drug-metabolizing ensemble was characterized by global proteomics based on the total protein approach [14]. The HLM samples in this series were selected to represent different degrees of alcohol exposure of the liver donors. It contained 8 HLM samples from non- and light drinkers (<=1 drink per day), 8 samples from moderate drinkers (2-3 drinks per day), and 7 samples from heavy drinkers (>=4 drinks per day). The results of proteomic characterization of HLM preparations may be found in the dataset available at Mendeley Data [15].

A series of titration curves of MONA demethylation obtained with a series of 23 individual HLM preparations is shown in Fig. 5a. All SSPs are normalized on the concentration of CPR in the samples. As seen from this figure, the probed preparations differ considerably in both the shape and the amplitude of the kinetic traces. To examine the correlations of their kinetic parameters with the level of alcohol exposure and the composition of the P450 pool, we subjected this dataset to Principal Component Analysis (PCA) and PCA-based global fitting. The main panel in Fig. 5b shows the first two principal components obtained with PCA. These components cover 99.5% of the differences between the traces. Assuming that these PCs represent different combinations of the same set of two Michaelis-Menten dependencies, we used a combination of a linear multidimensional regression algorithm and the Neelder-Mead nonlinear regression technique (see Materials and Methods) to find their parameters. This optimization yielded Michaelis-Menten dependencies with *K*_m_ values of 11 and 553 µM (Fig. 5b, insert) whose linear combinations approximate the first and second PC vectors with square correlation coefficients (*R*^2^) of 0.9995 and 0.997, respectively. Approximations of the individual SSPs in the set with these two Michaelis-Menten components are characterized by *R*^2^ values >0.975 for all curves, reaching >0.992 for 18 of them.

**Figure 5.**
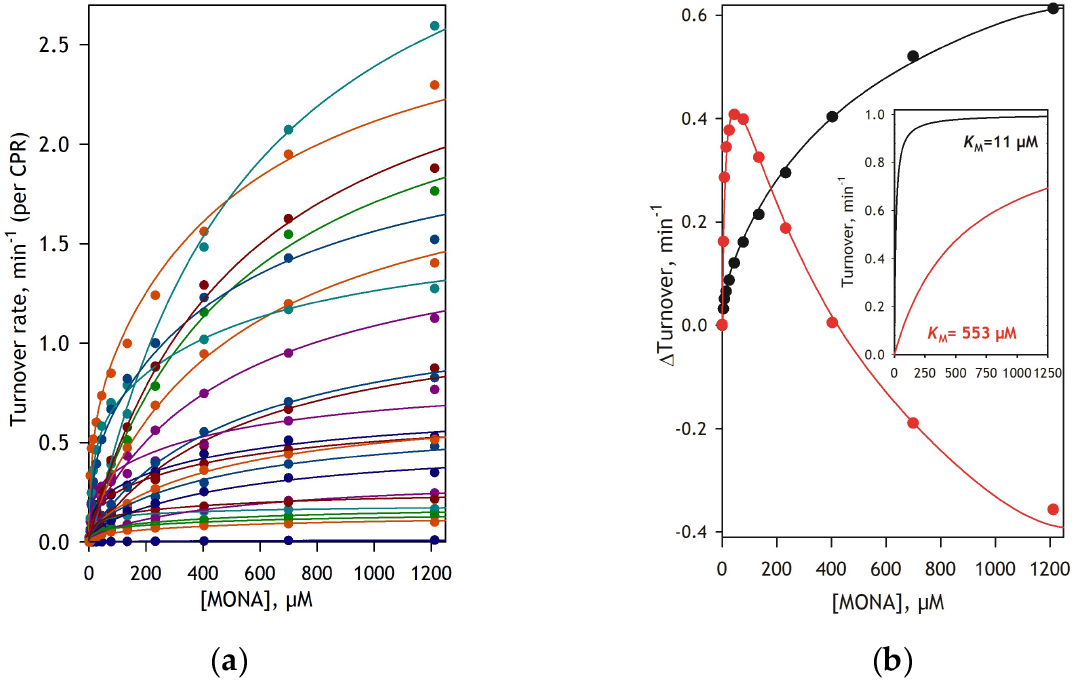
Substrate saturation profiles of MONA metabolism by 22 HLM preparations and their global analysis. Panel **(a)** shows the SSPs obtained with 22 HLM samples along with their approximations (lines) obtained from global fitting of the dataset with a combination of two Michaelis-Menten equations. Panel **(b)** shows the plots of the first (black) and the second (red) principal components found by PCA applied to the shown dataset. Together, these two principal components account for 99.5% of the overall differences among the SSPs. The inset in this panel shows two Michaelis-Menten components obtained from the global fitting of the dataset and used to approximate the principal components and the SSP traces shown in this panel.

The predominating (>50% of the total *V*_max_ for 19 out of 22 SSPs) low-affinity (*K*_M_=553 µM) component undoubtedly represents the activity of CYP1A2 (*K*_M_=591 ± 101, see Table 1). Although the origin of the minor high-affinity (*K*_m_=11 µM) component is uncertain, it may reflect the contribution of CYP2A6.

To probe the effect of alcohol consumption on MONA demethylation, we assessed the dependence of the maximal rates of the low-affinity (*V*_max1_) and the high-affinity (*V*_max2_) components on the Provisional Index of Alcohol Exposure (PIAE). This parameter was introduced in our recent study of the effect of alcohol exposure on the proteome of drug-metabolizing enzymes and transporters in the human liver [12]. PIAE is the combination of the relative abundances of four protein markers of alcohol exposure - the heat-shock 70 protein HSPA5 (GRP75), protein disulfide isomerases P4HB and PDIA3, and carboxylesterase CES2. A strict correlation of PIAE with the level of alcohol consumption reported by liver donors has been demonstrated in our previous study [12]. This index varies from ∼0 to ∼5, and its numeric value roughly approximates the average number of alcoholic drinks per day consumed by the liver donor.

Dependence of *V*_max1_ on PIAE is illustrated in Fig. 6A. As shown in this plot, the dataset as a whole does not reveal a significant correlation between CYP1A2 activity and PIAE. At the same time, the correlation of *V*max1 with the relative abundance (RA) of CYP1A2 is well pronounced. It is characterized by a correlation coefficient of 0.55 and a *p*-value of 8·10^-3^. However, it is well established that tobacco smoking results in largely increased CYP1A2 abundance in the liver [26]. The effect of smoking may interfere with the effect of alcohol exposure and mask its effect on CYP1A2 activity. The correlation between *V*_max1_ and CYP1A2 abundance in our dataset is largely driven by two HLM samples with abnormally high CYP1A2 levels from heavy smokers (>2 packs/day; donors HMC765 and HMC750). Excluding these two samples along with the two with the lowest PIAE values (<0.3, HMC 714 and HMC808) and the sample from the heaviest drinker in the pool (HMC553, PIAE=4.53) exposes a significant increase in CYP1A2 activity at increasing PIAE (black straight line in Fig. 6a, *R*=0.73, *p*=6.5·10^-4^) despite decreased CYP1A2 abundance in heavy and moderate alcohol consumers [12].

**Fig. 6.**
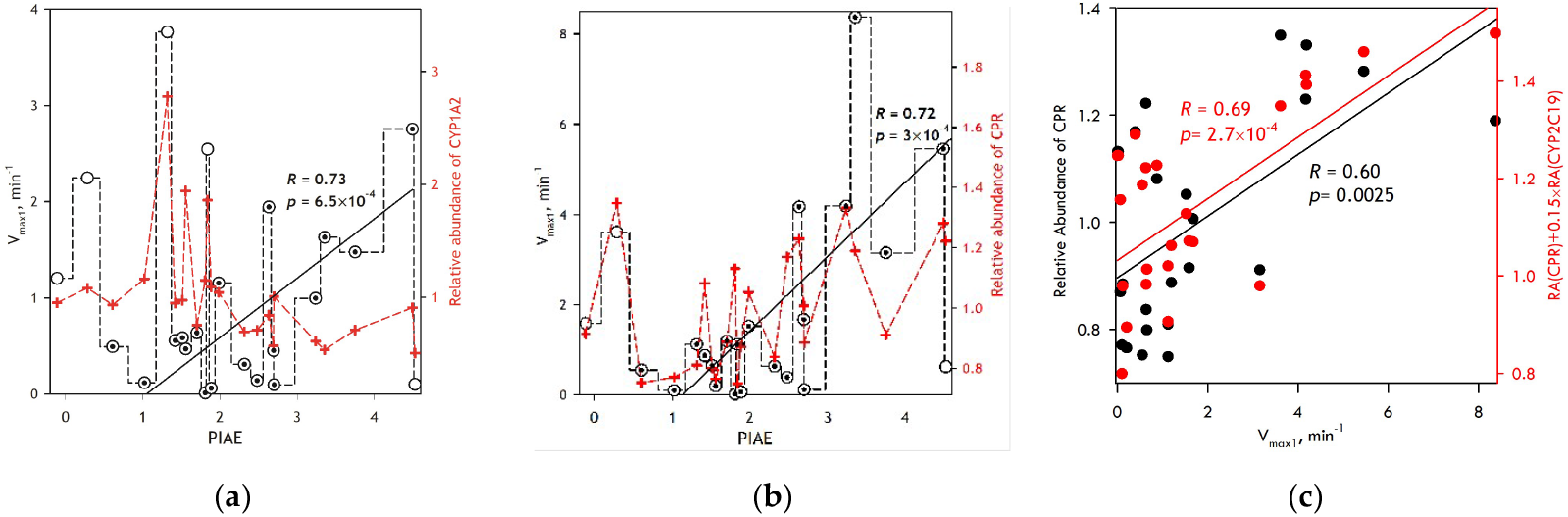
Correlations of the maximal rate of the CYP1A2-dependent phase of MONA demethylation (*V*_max1_) with the degree of alcohol exposure and the composition of the cytochrome P450 pool in 23 HLM preparations. Panels **(a)** and **(b)** show *V*_max1_ values normalized on CPR (panel **(a)**) or CYP1A2 (panel **(b)**) content plotted versus PIAE (black dotted and undotted circles and stepped dashed lines). Black, solid straight lines illustrate the correlation between Vmax1 and PIAE in a subset of HLM samples identified by dotted circles. Red crosses and red dashed lines show the dependence of CYP1A2 relative abundance on PIAE. Panel **(b)** illustrates the correlations of *V*_max1_ normalized on CYP1A2 content with the profile of relative abundance of CPR (black) and its combination with the RA profile of CYP2C19 (red).

A more thorough analysis of this data led us to the conclusion that the approach we used to normalize the reaction rate in the HLM samples may not be optimal in this case. Our usual method of normalizing reaction rates is to express them relative to the concentration of NADPH-cytochrome P450 reductase, the limiting component of the multienzyme cytochrome P450 system. While this approach is perfectly justified for substrates metabolized by multiple P450 enzymes, it may be misleading when the measured rate reflects the activity of only one P450 species. Since the low-affinity metabolism of MONA is catalyzed exclusively by CYP1A2, it is deemed more appropriate in this case to normalize the reaction rate to the concentration of CYP1A2.

Although our proteomic analysis only determined protein relative abundances (RA, the ratio of protein content in each preparation to its average across all samples in the set), a reliable estimate of absolute abundance can be obtained by multiplying the RA value by the average protein content determined in pooled HLM samples. According to our analysis of 11 pooled HLM preparations using targeted proteomics [9, 10], the average CYP1A2 content in HLM is 31.6±2.0 pmol/mg protein (see the dataset available in). Using this value, we can estimate the approximate absolute abundance of CYP1A2 in our samples and calculate the reaction rate, expressed as moles of substrate per mole of CYP1A2 per minute.

The resulting dependence of *V*_max1_ on PIAE is shown in Fig. 6b. As seen from this plot, scaling the reaction rate in this way makes its correlation with the degree of alcohol exposure much better pronounced. Although the correlation coefficient for the whole dataset is not high (0.42), excluding the two samples with the lowest PIAE values and the sample from the heaviest drinker in the pool, which shows abnormally low P450 activity, yields a correlation coefficient of 0.72 (*p*=6.5·10^-4^).

Assessing the relationship between the hereby normalized reaction rate and the relative abundances of P450 species and their redox partners (CPR and cytochrome b5), we observed the strongest correlation with CPR abundance (*R*=0.60, *p*=2.5·10^-3^). This observation, illustrated in Fig 6c, suggests that the increase in CYP1A2 activity with increased alcohol exposure is, at least in part, driven by alcohol-induced increases in the abundance of CPR in HLM [12]. The best combination of two RA profiles comprises CPR and CYP2C19, with multiplication factors (MF) of 1 and 0.145, respectively. This combination increases the correlation coefficient to 0.69 and yields a *p*-value as low as 2.7·10^-4^.

The dependence of *V*_max2_ on PIAE is illustrated in Fig. 7. Probing its correlation with the RA profiles of the members of microsomal P450 ensemble yields CYP2E1 as the best match (*R*=0.68, *p*=3.3·10^-4^, Fig. 7a). Combining the CYP2E1 RA profile with that of CYP2A6 with the factor of 0.171 increases the correlation coefficient to 0.70 (*p*=1.9·10^-4^). Notably, since CYP2E1 does not metabolize MONA, this observation may suggest that the high-affinity component reflects the activity of CYP2A6 in the complex with CYP2E1. The best combination of three RA profiles also includes the CPR profile (*R*=0.76, *p*=2.5·10^-5^, Fig. 7b).

**Fig. 7.**
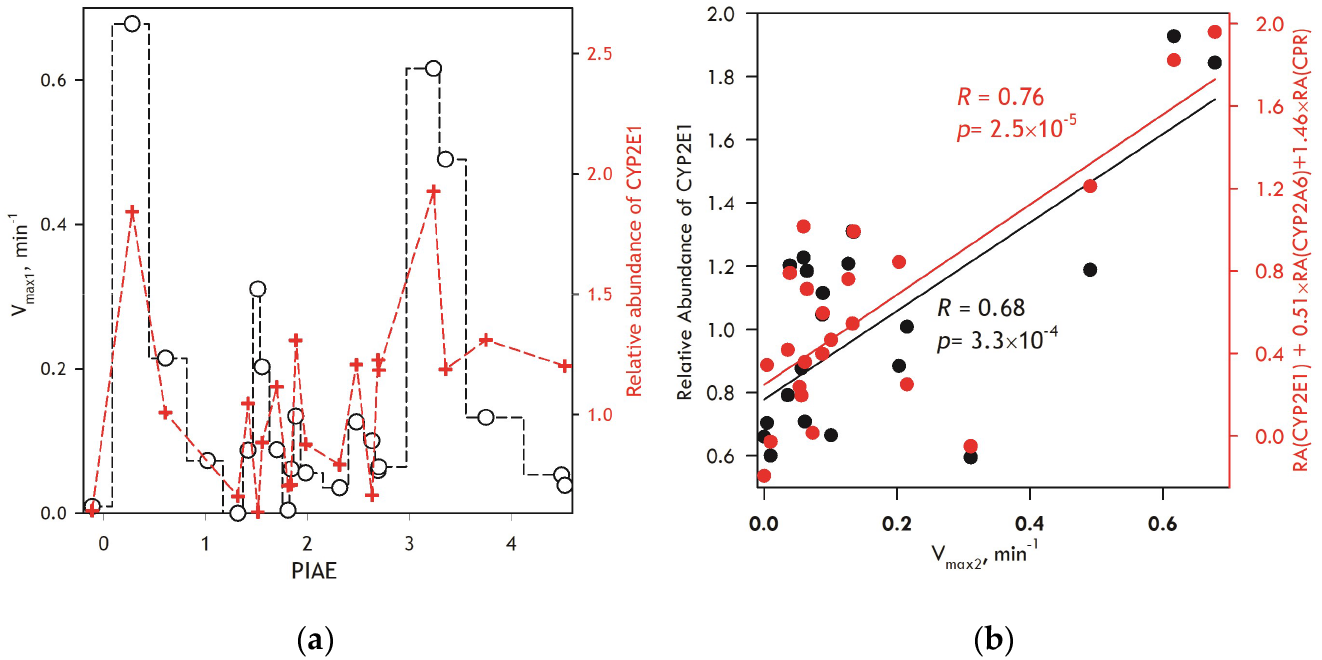
Correlations of the maximal rate of the high-affinity phase of MONA demethylation (*V*_max2_) with the degree of alcohol exposure and the composition of the cytochrome P450 pool in 23 HLM preparations. Panel **(a)** shows *V*_max2_ values normalized on CPR content plotted versus PIAE (black circles and stepped dashed line). Red crosses and red dashed lines show the dependence of CYP2E1 relative abundance on PIAE. Panel **(b)** illustrates the correlations of *V*_max2_ with the profile of relative abundance of CYP2E1 (black) and its combination with the RA profile of CYP2A6 and CPR (red).

### 3.4. Probing the effect of CYP2E1 on demethylation of MONA by CYP 2A6 and CYP1A2

According to the results of a global analysis of our dataset (Figure 5), the SSPs of MONA metabolism in human liver microsomes can be represented as a combination of two Michaelis-Menten equation components. Whereas the predominant low-affinity component undoubtedly reflects CYP1A2 activity, the nature of the second, high-affinity component remains uncertain. Although the correlation of its amplitude with the abundance of CYP2A6 suggests its attribution to the activity of CYP2A6, the *K*_m_ of 11 µM obtained from global analysis of the HLM dataset (Fig. 5) is considerably lower than that obtained with recombinant CYP2A6 in Bactosomes (*K*_M_≅35 µM), or CYP2A6-incorporated Supersomes (*K*_M_≅54 µM; see Table 1).

Based on the strong correlation between the amplitude of this high-affinity component and the CYP2E1 content, we hypothesized that the interaction between CYP2E1 and CYP2A6 proteins may lead to an increase in the affinity of CYP2A6 to MONA. To probe this hypothesis, we co-incorporated purified CYP2E1 and CYP2A6 proteins in a 1:1 molar ratio into Supersomes containing CPR and cytochrome *b*_5_ proteins, but with no P450 co-expressed. As shown in Fig. 8a, co-incorporation of CYP2E1 along with CYP2A6 results in a noticeable increase in CYP2A6 affinity for MONA but have no significant effect on the *V*_max_ value. Comparing the parameters of MONA metabolism by this preparation with that of CYP2A6-incorporated Supersomes, we found that the presence of CYP2E1 in the membrane causes a decrease in CYP2A6 *K*_M_ for MONA from 44.2 to 12.8 µM (Table 2), making it close to the value of 11 µM obtained from our global analysis of HLM dataset (Fig. 5). This result confirms our suggestion that the high-affinity component represent the activity of CYP2A6 enzyme in the complex with CYP2E1.

**Table 2.** Parameters of MONA demethylation by recombinant CYP1A2 and CYP2A6 enzymes after incorporation of recombinant CYP2E1.*

| Enzyme | $K_M$, $\mu\text{M}$ | $V_{max}$, $\text{min}^{-1}$ |
| --- | --- | --- |
| CYP1A2 in Supersomes (Control) | $655 \pm 132$ | $12.1 \pm 3.9$ |
| CYP1A2 + CYP2E1-incorporated Supersomes | $672 \pm 39$ | $19.3 \pm 0.7$ |
| CYP2A6-incorporated Supersomes (Control) | $44.2 \pm 12.2$ | $0.33 \pm 0.13$ |
| CYP2A6 + CYP2E1-incorporated Supersomes | $12.8 \pm 3.5$ | $0.30 \pm 0.05$ |

\* The values shown in the table represent the averages of the results of fitting of 2 (CYP1A2 Control), 2 (CYP1A2 + CYP2E1-incorporated Supersomes), 8 (CYP2A6 Control), and 3 (CYP2A6 + CYP2E1-incorporated Supersomes) individual datasets to the Michaelis-Menten equation. The ± values are the confidence intervals calculated for $p=0.05$ .

**Figure 8.**
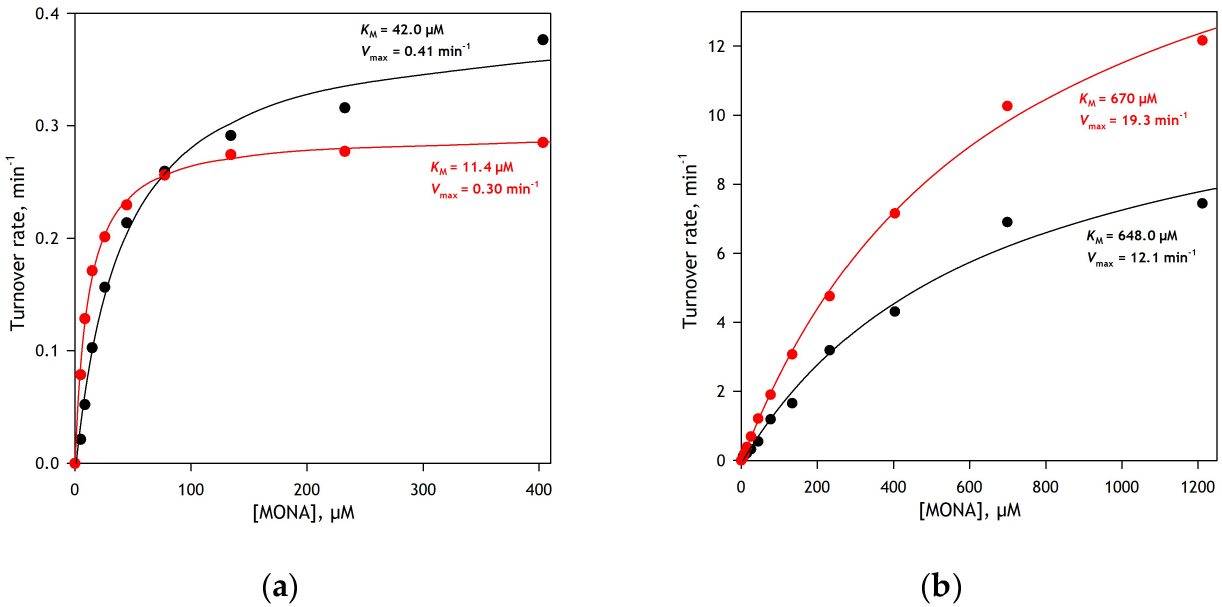
Effect of co-incorporation of purified CYP2E1 into Supersomes containing CYP2A6 (panel (***a***)) and CYP1A2 (panel (***b***)) on Substrate Saturation Profiles of MONA O-demethilation. The datasets represent the averages of the results of 3 (CYP2A6) and 2 (CYP1A2) individual experiments. The profiles obtained with CYP2E1 co-incorporated are shown in red. Solid lines show the approximation of the data sets with the Michaelis-Menten equation.

We previously demonstrated that the rate of CYP1A2-dependent metabolism of a fluorogenic substrate, 3-Cyano-7-ethoxycoumarin (CEC), was considerably increased after enriching the microsomes with CYP2E1 protein. This observation was interpreted as an indication of activation of CYP1A2 by its interactions with CYP2E1 [8]. Although our analysis did not reveal a correlation between the rate of MONA metabolism and the abundance of CYP2E1, we decided to test whether the interactions with CYP2E1 might increase CYP1A2 activity in MONA demethylation in a direct experiment. We incorporated purified CYP2E1 into CYP1A2-containing Supersomes and probed its impact on their activity. As illustrated in Fig. 9b and Table 2, we observed a significant increase in *V*_max_ with no change in *K*_M_. These results confirm our findings that interactions between CYP2E1 and CYP1A2 result in substantial activation of the latter.

## 4. Discussion

In this study, we introduce MONA as a new fluorogenic substrate with unique specificity to CYP1A2. The only other P450 enzyme capable of MONA O-Demethylation is CYP2A6. However, according to our study with recombinant enzymes in Supersomes, the *V*_max_ exhibited by CYP2A6 is over 25 times lower than that of CYP1A2. Most importantly, MONA is not metabolized by CYP1A1 and CYP2C enzymes, which distinguishes it from all other known fluorogenic CYP1A2 substrates.

Given the great superiority of CYP1A2 over CYP2A6 in the rate of MONA metabolism, and considering the comparable abundance of both enzymes in HLM, it was quite unexpected to observe biphasic substrate saturation profiles with a high-affinity (apparently CYP2A6-dependent) phase accounting for up to 60% of the total amplitude in some HLM samples. Another unexpected observation was the *K*_M_ value for this component, which was as low as 11 µM, much lower than the 54 µM observed with recombinant CYP2A6 (Table 1).

Close correlation of the amplitude of this phase with combined RA profiles of CYP2E1, CYP2A6, and CPR (Fig. 6c) suggests that it may represent the activity of CYP2A6 in the complex with CYP2E1, where its affinity to MONA increases. This hypothesis was confirmed by the experiment with co-incorporation of CYP2E1 into CYP2A6-containing Supersomes (Fig. 8a). However, the high fractional amplitude of this component in some HLM preparations lacks an apparent explanation, as interactions with CYP2E1 do not appear to increase the CYP2A6 turnover rate.

A clue to this discrepancy may be provided by low CYP1A2 activity in most of the analyzed HLM samples. As seen in Fig. 6b, when the *V*_max_ value of the CYP1A2-dependent low-affinity component is normalized to CYP1A2 content, it does not exceed 2 min^-1^ for most samples from donors with PIAE<3 (i.e., all donors except heavy drinkers). At the same time, the turnover number exhibited by the recombinant enzyme in Supersomes is as high as 11.9 min^-1^. Notably, the absolute value of *V*_max_ of the high-affinity component (*V*_max2_) does not exceed 0.7 min^-1^, which is commensurate with the rate of CYP2A6 turnover in Supersomes (0.5 min^-1^, see Table 1). Therefore, the high fractional amplitude of the CYP2A6-dependent component of MONA metabolism in some HLM preparations is due to low CYP1A2 activity in these samples rather than being an indicator of increased CYP2A6 activity.

An unexpectedly low CYP1A2 activity in HLM was also observed in our studies of the effect of CYP2E1 on the metabolism of 7-ethoxy-4-cyanocoumarin (CEC), the substrate metabolized by both CYP1A2 and CYP2C19 [8]. In that study, we demonstrated that, despite the much higher abundance of CYP1A2 in HLM and its higher turnover rate in CEC metabolism compared to CYP2C19, the SSPs of CEC deethylathion by pooled HLM samples are consistent with a predominant role of CYP2C19. We also found that enrichment of HLM with CYP2E1 causes an increase in CYP1A2 involvement in CEC metabolism that goes together with an increase in the rate of dealkylation of CYP1A-specific substrate 7-ethoxyresorufin [8]. Activation of CYP1A2 by its interactions with CYP2E1 was also demonstrated in the present study using MONA as a substrate (see Fig. 8 and Table 2).

However, what could be the reason for such low CYP1A2 activity in HLM, and what is the mechanism of its activation in alcohol consumers? Our understanding of these observations is based on the hypothesis of “positional heterogeneity” in P450 oligomers in the membrane. According to this hypothesis, P450 subunits that form homo- and heterooligomers in the membrane are not identical in their conformation and orientation. They differ in their abilities to be reduced, bind substrates, and interact with redox partners. In fact, oligomerization of cytochromes P450 results in the abstraction of a significant portion of the P450 pool from catalytic activity. From the standpoint of this hypothesis, the most obvious explanation for the low activity of some P450 species in HLM, noncommensurate with their high abundance, might be a difference in the propensities of different P450 species to occupy positions in the oligomer [27]. In particular, it might be hypothesized that, at the “average” composition of the P450 pool in HLM, the predominant part of CYP1A2 occupies obstructed positions in heteromeric complexes with other abundant P450 species, such as CYP3A4, CYP2C9, or CYP2A6. However, when CYP1A2 interacts with alcohol-inducible CYP2E1, the former preferentially occupies the “active” positions, whereas CYP2E1 fills the “obstructed” ones. This redistribution increases the active fraction of CYP1A2 in HLM.

Another important observation made in this study concerns the effect of CPR concentration on the activity of CYP1A2 and CYP2A6. As illustrated in Fig. 6c, the increased activity of CYP1A2 in alcohol consumers is largely due to an alcohol-induced increase in CPR content. Similar observation is true for CYP2A6 as well – the best correlation of the rate of CYP2A6-dependent component of MONA demethylation is exhibited by a combination of relative abundances of CYP2E1, CPR and CYP2A6 (Fig. 7b). It is to note, that the concentration of CPR in most of the samples in our set, regardless of the level of alcohol consumption, is higher that those of either CYP1A2 or CYP2A6. Thus, the effect of CPR on enzymes’ activity could be attributed to the allosteric effects of their interactions with CPR [28, 29, 30] rather than to the impact of their competition with other P450 species for limiting CPR. A possible mechanism of this effect may involve dissociation of P450 oligomers promoted by CYP1A2-CPR or CYP2A6-CPR interactions.

Regardless of the mechanistic basis of the activating effect of alcohol-induced expression of CPR and CYP2E1 on the activity of CYP1A2, it has a critical importance for practical pharmacology as it affects the pharmacokinetics of several antidepressants and antipsychotics metabolized by CYP1A2, including those used for the treatment of alcohol use disorders (AUD) and alcohol withdrawal syndrome (AWS). The alcohol-induced increase in CYP2A6 activity suggested by our study may play a role in the pharmacological interactions between smoking cessation and alcohol consumption.

## Author Contributions

Conceptualization, D.R.D.; methodology, D.R.D., B.P.; software, D.R.D.; formal analysis, K.G., D.R.D.; investigation, K.G., N.D., K.P., D.K.S.; data curation, D.R.D., K.G., D.K.S.; writing—K.G., D.R.D.; writing—review and editing, D.R.D.; supervision, D.R.D.; funding acquisition, D.R.D., B.P. All authors have read and agreed to the published version of the manuscript.

## Institutional Review Board Statement

Ethical review and approval were waived for this study because it was performed with IRB-exempt deidentified tissue specimens.

## Informed Consent Statement

Not applicable.

